# Host community structure can shape pathogen outbreak dynamics through a phylogenetic dilution effect

**DOI:** 10.1101/2023.05.23.541907

**Authors:** Marjolein E.M. Toorians, Isabel M. Smallegange, T. Jonathan Davies

## Abstract

Biodiversity loss and anthropogenic modifications to species communities are impacting the frequency and magnitude of disease emergence events. These changes may be related through mechanisms in which biodiversity either increases (amplifies) or decreases (dilutes) disease prevalence. Biodiversity effects can be direct, when contacts among competent hosts are substituted by contacts with sink hosts, or indirect through regulation of host abundances. Whether a disease is diluted or amplified depends on the competence of the host species in the community. Here, we introduce a multi-host compartmental disease model in which we assume that host competence is determined by evolutionary relatedness. By simulating communities of varying phylogenetic structure, and estimating community disease outbreak potential (*R*0), we show how differences in host phylogenetic relatedness can switch host communities from diluting to amplifying a disease, even when species richness is unchanged. We additionally show that phylogenetic dilution can occur simultaneously with amplification through species richness, highlighting the multidimensionality of the disease-diversity relationship. We illustrate our model using empirical data describing the relationship between phylogenetic distances separating hosts and their likelihood of disease sharing. Our study demonstrates how a phylogenetic dilution effect can emerge when we allow transmission to depend on host evolutionary histories.

## 1 Introduction

Over the past century, anthropogenic pressures on the environment have accelerated (Steffen et al., 2015), leaving many species threatened with extinction (IPBES, 2020; Barrett et al., 2018; IUCN, 2021). Coinciding with increasing in biodiversity loss has been an increase in the frequency of disease outbreaks (Daszak et al., 2001; UN, 2020) and it has been suggested that these two trends may be linked (Halliday and Rohr, 2019; Rohr et al., 2020). Similar patterns are seen for plants and animals (Daszak et al., 2000; Woolhouse et al., 2005; Jones et al., 2008), suggesting a common process. However, there remains debate about how shifts in the composition and richness of host communities may translate to changes in disease prevalence and outbreak potential (LoGiudice et al., 2003; Joseph et al., 2013). While much attention has been given to the relationship between host species diversity and disease (e.g. Halliday and Rohr, 2019; Rohr et al., 2020; Keesing et al., 2006, 2010; Han et al., 2020), predicting how changes in host species composition may alter disease prevalence requires a mechanistic understanding of multi-host disease dynamics (Woolhouse et al., 2001; Dobson, 2004; Luis et al., 2018).

Theoretical models suggest that an increase in host species richness can either decrease disease prevalence (dilution effect: Table 1) or increase disease prevalence (amplification effect: Table 1) (Keesing et al., 2006, 2010; Ostfeld and Keesing, 2012). Both dilution and amplification can occur via changes in transmission, either directly through modifying encounter rates between competent and less-competent hosts or indirectly through changing densities of competent hosts (Rudolf and Antonovics, 2005; Ostfeld and Keesing, 2000). In practice, the distinction between the processes of amplification and dilution is not straightforward. Some diseases appear to be diluted by diversity in some instances, but amplified by diversity in other instances, depending on the host community species composition, and host shedding rates or relative abundances (Barasona et al., 2019; Streicker et al., 2013; Stewart Merrill et al., 2022). It is also possible for both dilution and amplification to occur simultaneously in the same system, for example, depending on the density and composition of the host community, as has been suggested in rodents infected by hantavirus (Luis et al., 2018) and for RNA viruses in honey and bumble bees, where species abundance and richness separately had amplifying and diluting effects, respectively (Cohen et al., 2022). Keesing and Ostfeld (2021), who first coined the ‘dilution effect’, acknowledge that the general effect of biodiversity on disease prevalence remains controversial.

**Table 1:**
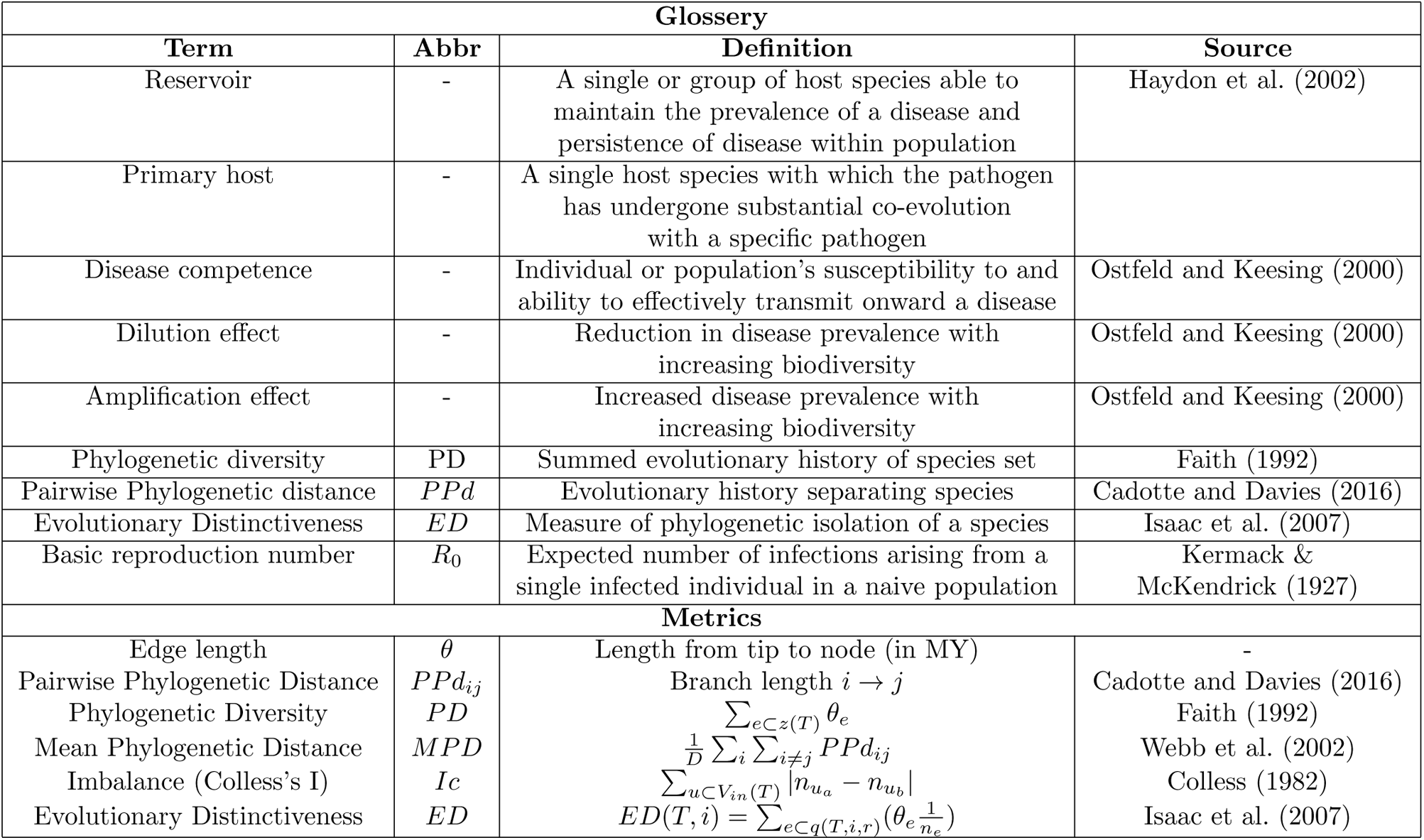
Glossary of terms and biodiversity metrics. For phylogenetic metrics, *θ_i_* is the branch length from species *i* to the root of the tree and *n* is the number of species in the tree. *e ⊂ z*(*T*) denotes the edges, *e*, in a set *z* for tree *T*, *D* is the total number of species combinations, and the set *q*(*T, i, r*) connecting species *i* to the root *r*, and *n_e_* is the number of species that descend from edge *e*. *V_in_*(*T*) is the set of all inner vertices of the tree, and 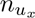 are the leaves in the subtree *x*.

Ultimately, it is the composition and relative competence of hosts within a community that drive dilution/amplification dynamics. Species that are able to maintain endemic prevalence—reservoir hosts (Table 1, Haydon et al., 2002)—have high competence, and can dominate transmission in a multi-host community. Less competent hosts, however, may function as dead-end hosts if they either do not maintain the disease, or do not contribute to onward transmission, thereby reducing disease spread. Data on host competence would allow us to make more accurate predictions about how changes in host diversity would impact disease prevalence; however, such data are frequently lacking, especially for low-competent hosts and hosts with no prior exposure. One approach to bridge this data gap is to approximate host competence from life history data and species’ physiology, such as body size or other life-history traits (Dobson, 2004; Joseph et al., 2013; Cohen et al., 2022). Here, we suggest that if relevant traits are evolutionarily conserved, species phylogeny might provide another useful and more general proxy.

There is a growing weight of evidence showing that diseases are more frequently shared between closely related hosts (Gilbert and Webb, 2007; Davies and Pedersen, 2008; Streicker et al., 2010; Parker et al., 2015; Farrell and Davies, 2019a). This phylogenetic structure in the distribution of pathogens among hosts likely reflects phylogenetic conservatism in evolved defense mechanisms (Streicker et al., 2010; Mollentze and Streicker, 2020). We might therefore expect the evolutionary distance between hosts to be correlated with both the probability of sharing a disease and within-host pathogen load following infection. For example, in grassland communities, disease prevalence was higher in species with many close relatives in the community (Parker et al., 2015), whereas hosts that were phylogenetically isolated (Table 1) in their communities obtained a *rare-species advantage* and were less likely to share pathogens with co-occurring species. Phylogenetic isolation—Evolutionary Distinctness (ED, Table 1)—from co-occurring hosts might, therefore, be an indicator of low competence for diseases most prevalent within a given host community (although they may be highly competent hosts for other diseases). We suggest that the phylogenetic composition of hosts may thus be an additional component in the establishment probability and disease dynamics of a pathogen in the community.

Here, using simulations and a simple compartmental (Susceptible-Infected) disease model, we explore the importance of the phylogenetic structure of host communities on outbreak potential (*R*_0_) for a multi-host disease, assuming density dependent (DD) dynamics, where the contact rate is constant. In Appendix Section 6 we show results for a frequency-dependent (FD) model. For simplicity, our model assumes no recovery, such as is the case for *Mycobacterium bovis* infection, the causative agent of bovine Tuberculosis (bTB) (Jolles et al., 2005; Ezenwa et al., 2010), but it could easily be extended. Scaling transmission probability in proportion to the inverse of the phylogenetic distance between hosts, such that transmission is more likely between closely related hosts, we consider two alternative scenarios: 1) transmission is proportional to the evolutionary distance between infected and susceptible hosts; and 2) transmission is proportional to the distance between the receiving (secondary) and primary (co-evolved) host, irrespective of the identity of the donating host (Figure 1). We provide examples of our model predictions parameterised using empirical data on plant pests (Gilbert et al., 2012). We suggest that our model may help in our understanding of the conflicting evidence for dilution and amplification effects of biodiversity across different host communities and disease systems.

**Figure 1:**
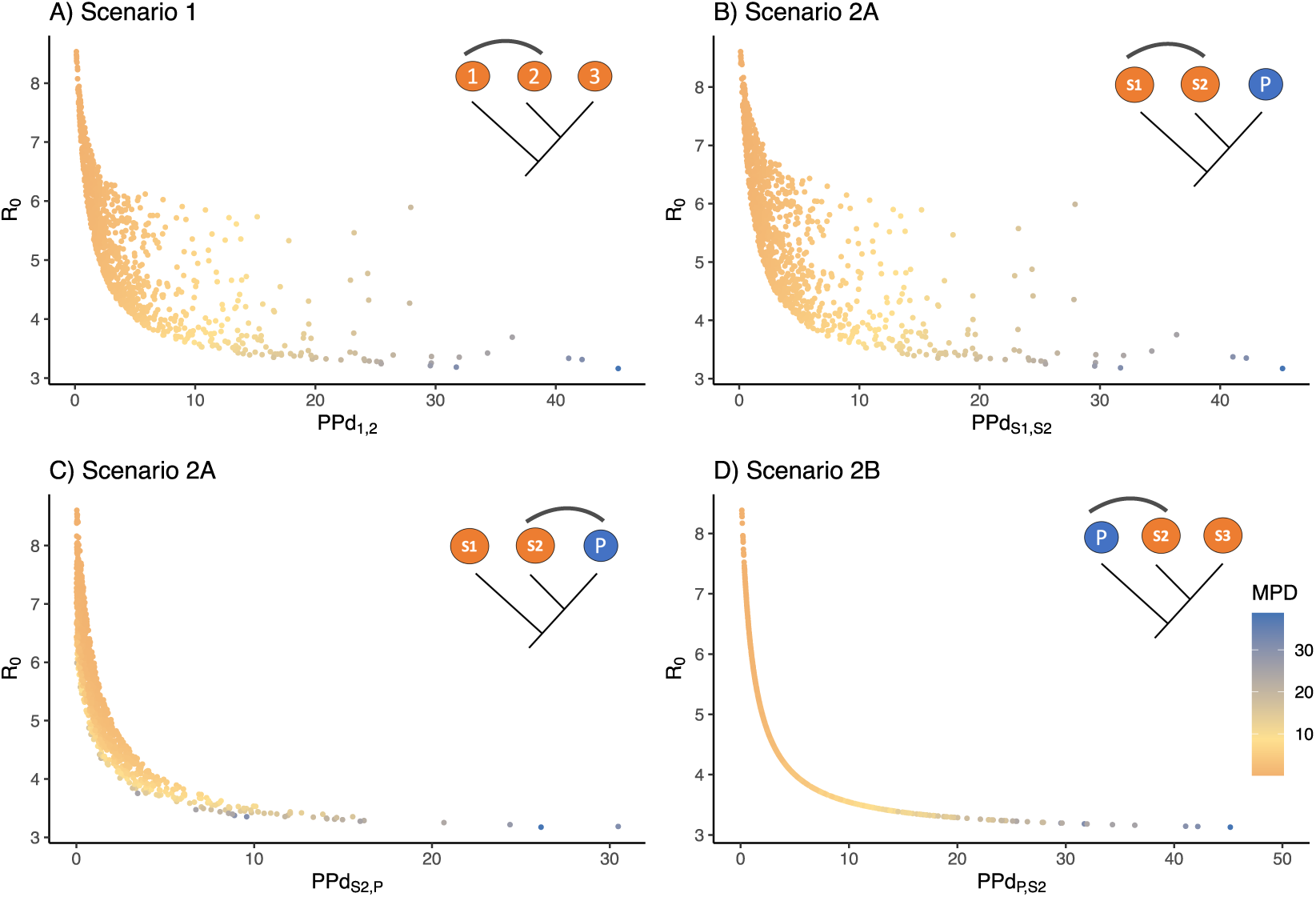
Community disease outbreak potential (*R*_0_) plotted against the phylogenetic distance (*PPd*) between host pairs indicated on the x-axes. Shading is proportional to *MPD*. Inset within each graph provides an illustration of the alternative primary-secondary host dynamics for a simple three-taxon host system. The first panel (A) shows scenario 1, with no assigned primary host. The second (B) and third (C) panels show scenario 2a (primary host as ingroup) with blue circles representing the primary host (P), and orange circles the secondary hosts (S1 and S2). (B) shows the pairwise phylogenetic distance (*PPd*) between the two secondary hosts (S1 and S2) on the x-axis, whereas (C) shows the *PPd* between the primary host (P) and secondary host 2 (S2). The fourth panel (D) shows Scenario 2b, with the primary host sister to both secondary hosts (primary host as outgroup). Here both secondary hosts are equidistant from the primary host, and thus the system converges on a simple two-host model, and *R*_0_ is effectively determined by tree depth. Grey line in illustration shows the distance between the two hosts on x-axis of each graph. Simulated data from the model with *α* = 1, *ν* = 0.1, *µ* = 0.2, *ψ* = 0.01.

## 2 Methods

The outbreak potential of a disease may be quantified by its reproductive number (*R*_0_), which represents the average number of new infections that can arise from a single infected individual over the course of their lifetime, considering a population of susceptible individuals. When this value is greater than one, an epidemic can occur. The *R*_0_ can be calculated from the ordinary differential equations describing the susceptible (*S*) and infected (*I*) individuals (Kermack and McKendrick, 1927; May and Anderson, 1984; Dobson, 2004):

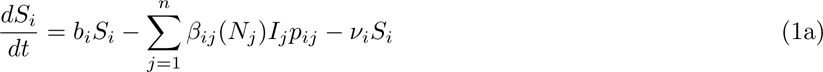

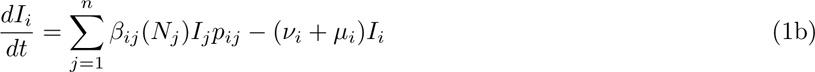

where *β*(*N_j_*) describes the transition from *S* to *I* compartments per time unit, *ν_i_* and *µ_i_* are the species-specific disease-induced and natural mortality rates, respectively, *b_i_* is the per-capita host birth rate, and *p_ij_* represents the mode of transmission, such that 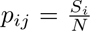 for frequency dependence, or 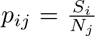 for density-dependence. We focus primarily on density-dependent transmission, as this is the most commonly assumed mode of transmission for wildlife diseases White et al. (2017), but provide matching results for frequency dependent dynamics in the supplement. Details of the model and its derivation can be found in Appendix Section 1.

Following Keeling and Rohani (2011), we can decompose the transmission rate parameter, *β* as:

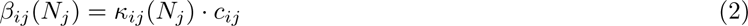

In this equation, the transmission rate, *β*, is a function of the probability of transmission given a contact, *c_ij_*, multiplied by the contact rate, *κ*. We can define *κ* as a function of the individuals in the community, *N_j_*, allowing this model to describe both frequency-dependent and density-dependent disease transmission, such that *κ* is either a constant or varies with *N_j_*, respectively (see Appendix Section 1). This dynamic for *κ* is captured in the Kronecker function (see Appendix section 4).

In a multi-host community, we can calculate *R*_0,community_ by using the Next-Generation-Matrix (Diekmann and Kretzschmar, 1991; Diekmann et al., 2010). A detailed derivation can be found in Appendix Section 1. In a 3-host system, matrix **K** can be represented as:

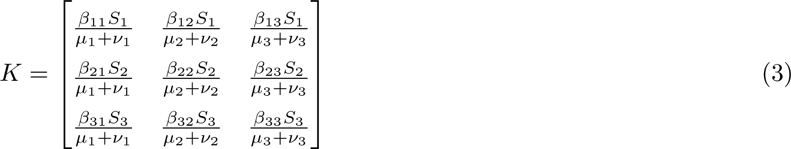

The maximum real eigenvalue of matrix **K** is the *community R*_0_ and describes the reproductive success of a generalist pathogen in the fully susceptible host community. On the diagonal of matrix **K** we can find the individual *R*_0_ of species *i* in isolation, *R*_0*,i*_. When *R*_0*,i*_ *>* 1, host species *i* is a reservoir host; when *R*_0*,i*_ *<* 1, host species *i* cannot sustain infection on its own, resulting in obligate multi-host parasitism (Fenton et al., 2015). The sum of a column of matrix **K** shows the total received infection of the *n^th^* species in the community (including intraspecific transmission). The sum of a row shows the total contribution of that species to the community infected class.

We can easily rescale the interspecific values for *c_ij_* in the *β* transmission rate term; here, we assume that probability of successful infection increases with decreasing evolutionary distance (Lively, 2010; Gilbert et al., 2012; Parker et al., 2015; Farrell and Davies, 2019a). Now, the probability of successful transmission from host *j* to *i* is determined by the evolutionary distance separating the two species (Pairwise Phylogenetic distance, *PPd*, see Table 1). While we model the effect of phylogenetic distance on the probability of transmission given a contact, we might expect contact rates to also co-vary with phylogeny. However, the weighting by phylogeny can be equally placed on either *κ* or *c*, and thus a single weighting term can capture phylogenetic structure in transmission probability, contact rate or both. We assume all species have equal abundances, thus allowing for encounter reduction with the addition of a species to the host community, putatively the main mechanism behind dilution (Keesing et al., 2006). We do not, therefore, explore potential effects of host evenness or other diversity indices combining richness with abundances, but it would be possible to further expand our model to consider more complex host population dynamics if there was evidence to support them in a given community. We use simulations to explore two alternative scenarios describing primary-secondary host dynamics.

### Scenario 1: No primary host

In this scenario, the probability of successful transmission from species *j* after contact with species *i*, is modelled as:

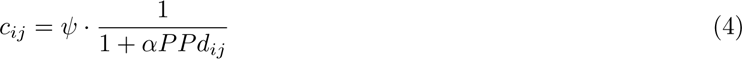

where *PPd_ij_* is the pairwise phylogenetic distance between hosts *i* and *j* (typically measured in millions of years), *α* is a scalar for *PPd*, and *ψ* scales the probability of infection. The probability of establishing an infection in a novel host simply depends on the distance between the transmitting, *j*, and receiving, *i*, host. For our simulations, we assume 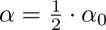 as *PPd* represents 2 * the time since divergence between hosts; however, alternative evolutionary models can be assumed that allow for non-linear relationships, or *α* can be estimated directly from empirical observations, as we describe below. This scenario might apply to diseases such as Rabies virus (*Rabies lyssavirus*), a multi-host pathogen affecting a large range of mammal species, where isolation within a species and high mutation rates cause rapid differentiation and local adaptation of the virus to its host (Bourhy et al., 1999; Streicker et al., 2010; Mollentze et al., 2014).

### Scenario 2: Single primary host

Here, we scale the interspecific probability of successful transmission by the distance between the secondary, recipient host (*i*) and the primary host, regardless of the identity of the proximate transmitting host. In this scenario, we assume the pathogen is best adapted to the primary host *P*, and so the probability of disease transmission will decreases with the evolutionary distance between the recipient host and the primary host, such that:

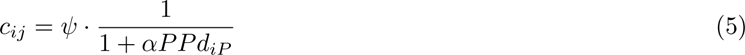

In a three-species system, there are two unique phylogenetic placements of the primary host, nested within the ingroup sister pair, or as outgroup to the sister pair, we refer to these alternative placements as ingroup and outgroup, respectively. Both scenarios are illustrated in Figure 1.

One well-described empirical example of a disease systems with primary-secondary host dynamics is bovine tuberculosis (bTB). In sub-Saharan Africa, bTB is thought to be endemic in African Buffalo (*Syncerus caffer*), which function as a reservoir, and primary host (Bengis and Erasmus, 1988; Jolles et al., 2005; De Garine-Wichatitsky et al., 2013; Ezenwa and Jolles, 2015). However, bTB frequently spills over to secondary hosts, including other ungulates, where it may establish persistent infections, and, more rarely, to phylogenetically distant species, which act as dead-end hosts with no forward transmission (Michel et al., 2006).

### Simulating communities

We used simulations to describe the modelled relationship between host community phylogenetic structure and disease outbreak potential (*R*_0_). We first simulate host phylogenetic topologies under a pure-birth Yule process, using the R-package *phytools* (Revell, 2012), with birth rate, *b*, drawn from an exponential distribution to simulate *waiting times* (the branching times between speciation events) (See Appendix Section 2 for more information). Communities simulated with low *b* have an increased distance between branching events, and are thus composed of more distantly related species than communities simulated with high *b* (Appendix Figure 2).

**Figure 2:**
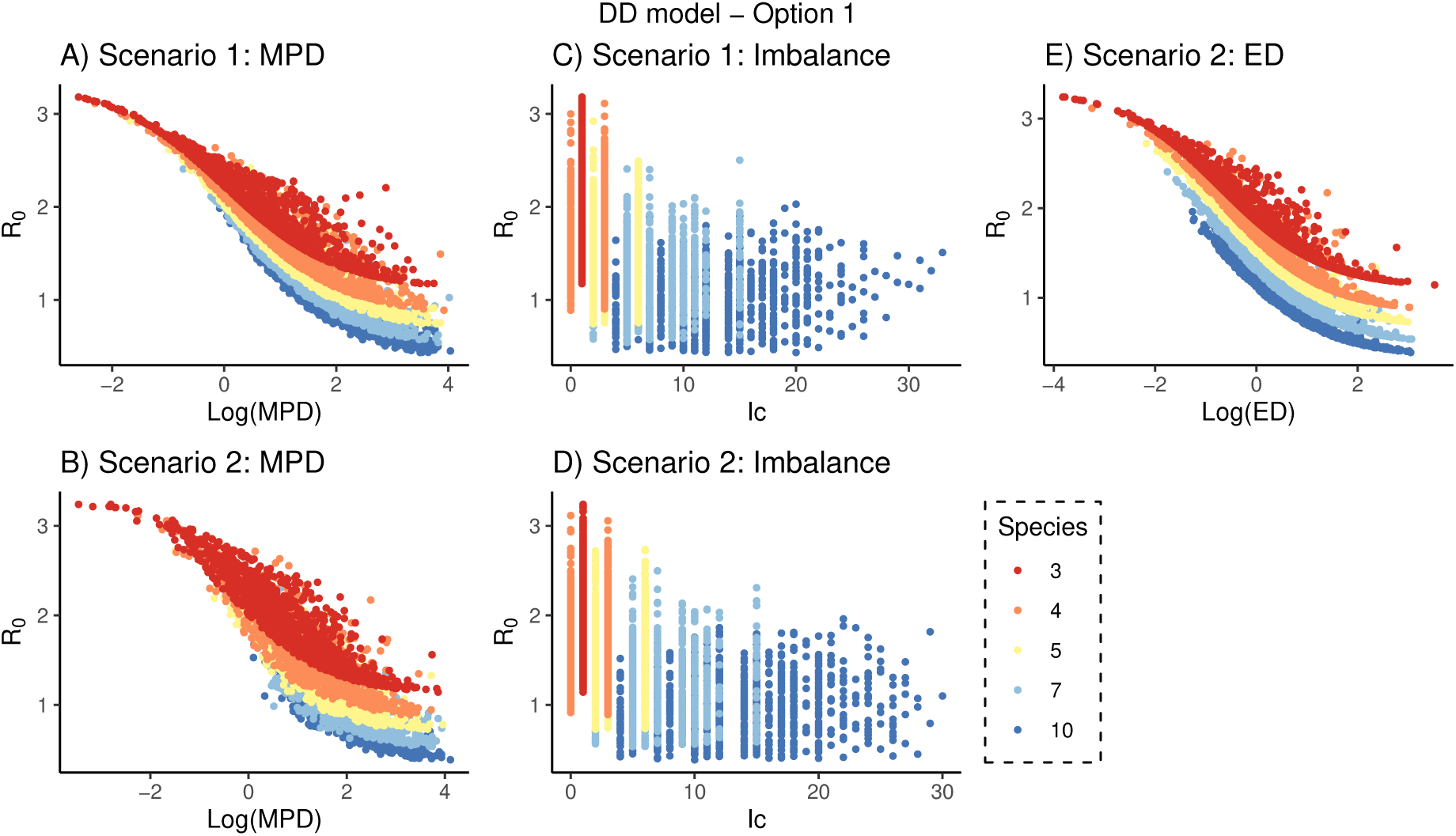
Modelled relationships between phylogenetic community structure and *R*_0_ assuming transmission success is inversely weighted by the phylogenetic distance between hosts and replacing individuals (Option 1, substitutive). Scenario 1 (no primary host): A) Mean Pairwise Distance (*MPD*) of the community is inversely related to *R*_0_. B) Relationship between *R*_0_ and the Mean Pairwise phylogenetic Distance (*MPD*) between host species, with outbreak potential decreasing logistically with increasing *MPD*. C) Imbalance of the phylogenetic tree representing the host species community shows a significant but weak trend for *R*_0_ to increase with increasing tree balance. Scenario 2 (single primary host). D) Relationship between *R*_0_ and phylogenetic tree Imbalance (*Ic*), shows high variation. E) Relationship between *R*_0_ and Evolutionary distinctiveness (ED) of the primary host, demonstrating similar dynamics to *MPD*. Data from simulations with *κ* = 1, *α* = 1, *ν* = 0.1, *µ* = 0.2 and *ψ* = 0.01. Colours indicate simulated communities with different host richness (*n*=3,4,5,7,10).

To study dilution and amplification effects, we explored two alternative effects on host densities, as described by (Ostfeld and Keesing, 2012): (1) Substitutive, adding a species to a community reduces the abundance of other species, such that the total number of individuals in the community, *N_T_*, is constant (where *N_T_* = *S_T_* + *I_T_*= 100, and *T* represents the total *n* species); (2) Additive, the total number of of individuals, *N*, increases when a novel species is introduced. Each species has the same number of individuals (*N_i_* = *N_j_* = *…* = *N_n_* =100). In all simulations *I ≈* 0 and *N ≈ S* for the calculation of *R*_0_.

We generated 100 host communities for different combinations of richness (*n*=3,4,5,7,10) and speciation rates (0 *< b <* 1). For each simulated community phylogeny, we calculated i) Phylogenetic Diversity (*PD*) to capture host tree size; ii) Mean Pairwise Distance between hosts (*MPD*) as a measure of phylogenetic community structure; and iii) Tree *Imbalance*, *Ic*, to capture the effect of tree-topology, (Table 1). To explore primary-secondary host dynamics in Scenario 2, for communities with *>* 3 hosts, we additionally recorded primary host Evolutionary Distinctiveness (*ED*), a metric describing the phylogenetic isolation of a tip on the phylogeny (Isaac et al., 2007). In simulations, the phylogenetic placement of the primary host (Scenario 2) was assigned randomly. We then calculated the community *R*_0_, with a natural and disease-induced mortality of *ν* = 0.1 and *µ* = 0.2, respectively and assuming *ψ* = 0.01 and *α* = 1 to return realistic values of *R*_0_ (0 *< R*_0_ *<* 3 for option 1 and 0 *< R*_0_ *<* 15 for option 2) (Roche et al., 2012). For results reported in the main text, we set contact rates, *κ_ij_* = 1, such that interspecific and intraspecific contacts are assumed equally likely. We explore alternative contact scenarios in Appendix section 4. We use regression models to quantify how different aspects of host phylogeny mediate disease outbreak potential. Both *MPD* and *ED* were log transformed, and all variables were scaled using a *z*-transform before model fitting. Colinearity among predictors was assessed using Variance Inflation Factors (VIF) (Zuur et al., 2010).

Finally, we parameterised our model using empirically derived transformations of the pairwise phylogenetic distances between hosts for viruses and bacteria from Gilbert et al. (2012). This study estimated host susceptibility in relation to the phylogenetic distance separating donating and recipient hosts for various pathogen groups using the Global Pest and Disease Database from the US Department of Agriculture Animal and Plant Health Inspection Services, a database containing 137,000 pest-host records for 18,500 plant species worldwide. Gilbert et al. (2012) assume the following functional relationship: *logit*(*c_ij_*) = *b*_0_ + *b*_1_ *∗ log*_10_(*PPd_ij_* + 1). Because the steepness of the slope describing the relationship between susceptibility and phylogenetic distance differed among pathogen types, we fit two separate models: *b*_0_ = 8.44 and *b*_1_ = −5.2, for viruses, and *b*_0_ = 3.26 and *b*_1_ = −2.97 for bacteria.

## 3 Results

We used simulations to explore the relationship between host community phylogenetic structure and disease outbreak potential assuming transmission success is a function of the phylogenetic distance between hosts. We show that both the phylogenetic structure of the host community and the phylogenetic placement of the primary host can have large influence on community *R*_0_.

### 3.1 Option 1: Substitutive

Adding a species to a community reduces the abundance of other species, such that the total number of individuals in the community remains constant.

#### Scenario 1: No primary host

In the absence of a single primary host, the establishment probability of the disease (*c*) simply depends on the phylogenetic distance between the receiving and donating hosts, as described in Eqn (4) and as illustrated in Figure 1A for the 3-taxon case. We modelled *R*_0_ as a function of *MPD*, species richness (*n*) and Imbalance (*Ic*), capturing the key axes of variation in our PCAs (Appendix Figures 3 and 4). There was strong covariation between *MPD* and *PD* (VIFs of 111 and 140, respectively), we report multivariate models including *MPD* here as it is independent of species richness, *n*; however, models exchanging *MPD* with *PD* were qualitatively similar.

**Figure 3:**
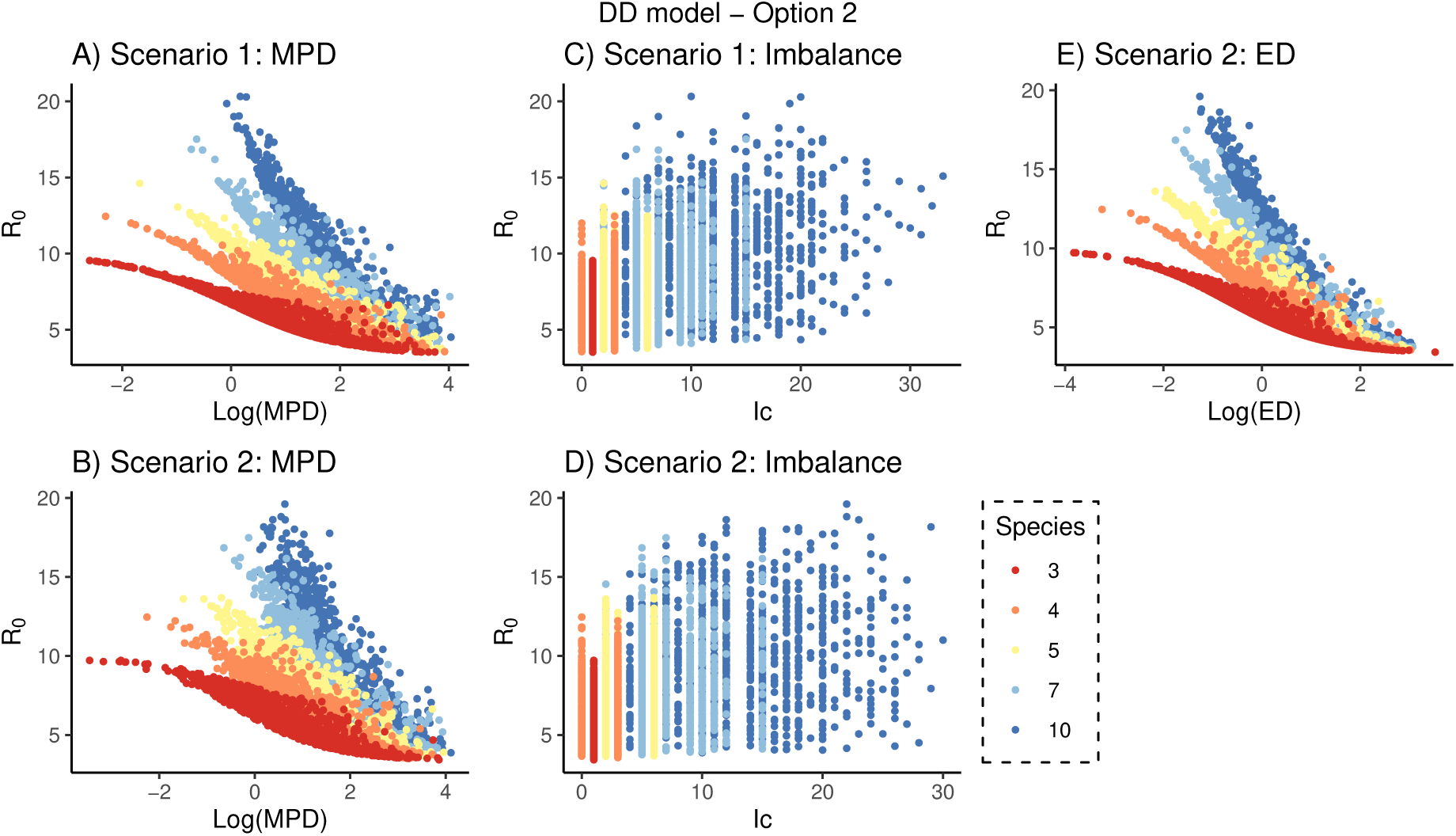
Modelled relationships between phylogenetic community structure and *R*_0_ assuming transmission success is inversely weighted by the phylogenetic distance between hosts and adding species increases total community size (Option 2, additive). Scenario 1 (no primary host): A) Mean Pairwise Distance (*MPD*) of the community is inversely related to *R*_0_. B) Relationship between *R*_0_ and the Mean Pairwise phylogenetic Distance (*MPD*) between host species, with outbreak potential decreasing logistically with increasing *MPD*, but here the relationship becomes less clear once species richness increases. C) Imbalance of the phylogenetic tree representing the host species community shows a significant but weak trend for *R*_0_ to increase with increasing tree balance. Scenario 2 (single primary host). D) Relationship between *R*_0_ and phylogenetic tree Imbalance (*Ic*), illustrating high variation. E) Relationship between *R*_0_ and Evolutionary distinctiveness (*ED*) of the primary host, demonstrating similar dynamics to *MPD*. Data from simulations with *κ* = 1, *α* = 1, *ν* = 0.1, *µ* = 0.2 and *ψ* = 0.01. Colours indicate simulated communities with different host richness (*n*=3,4,5,7,10).

**Figure 4:**
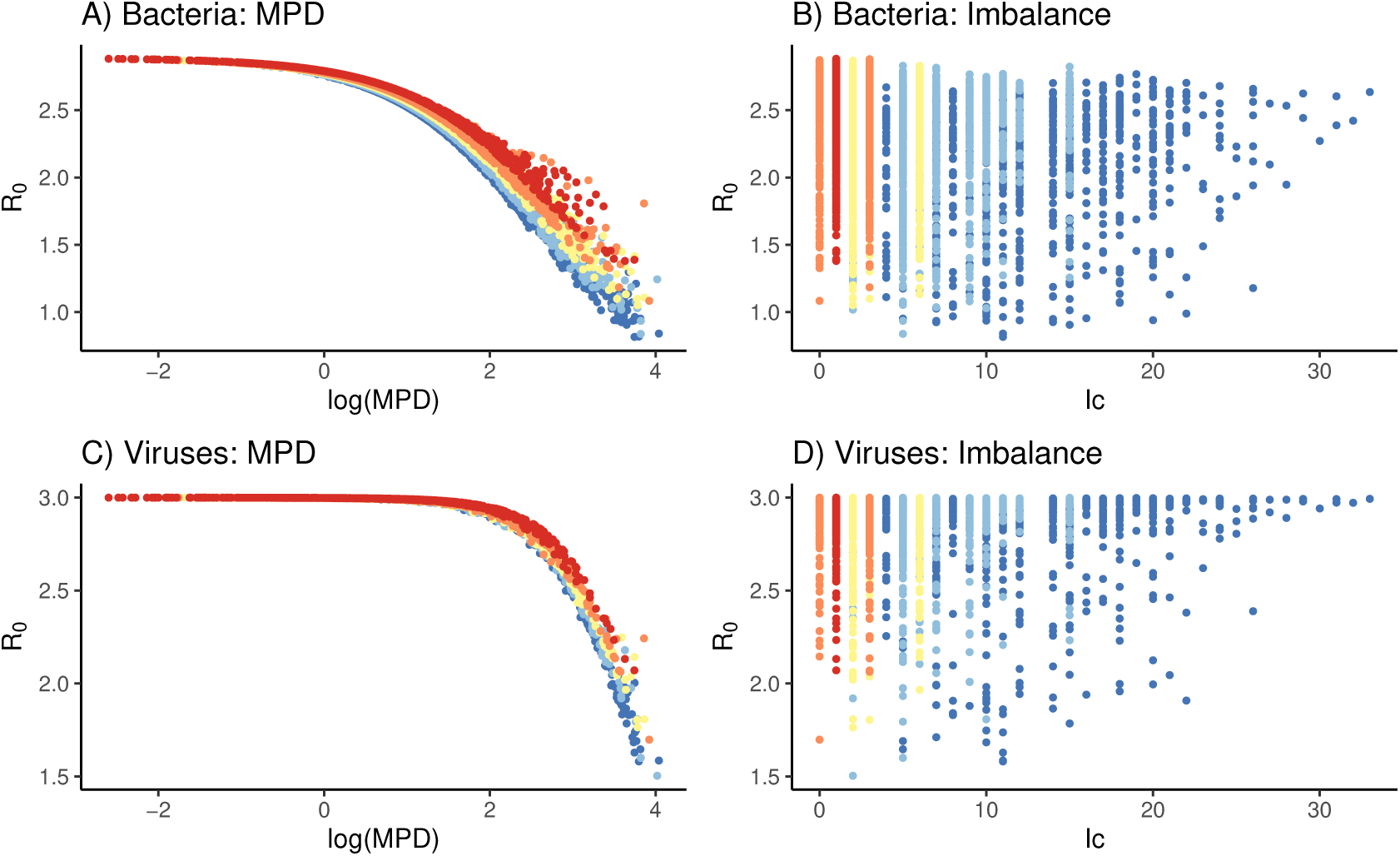
Modelled relationships between phylogenetic community structure and *R*_0_ assuming the phylogenetic transformation from Gilbert *et al*. (2012) for bacteria and viruses, Scenario 1 and model Option 1 (replacing individuals). A and C illustrate the effect of Mean Pairwise distance between hosts, B and D illustrate the effect of tree Imbalance.

In the single-predictor models (Table 2), *PD* explained the most variation in *R*_0_; more phylogenetically diverse communities generally had lower disease outbreak potential. *MPD* and host richness, *n*, were also strong predictors of *R*_0_ (Table 2), with communities composed of more, but less closely related species having a lower outbreak potential (Figure 2A). While Imbalance (Ic) was positively correlated with *R*_0_, more balanced trees had a higher outbreak potential, the relationship was weak (Figure 2C). In our multivariate model, *MPD* was the strongest predictor of *R*_0_ (Table 2), but both *n* and *Ic* explained some additional variation.

**Table 2:** Models describing the variation in *R*_0_ with various phylogenetic metrics for simulations with model Option 1 (replacing abundance) and *ν* = 0.1, *µ* = 0.2, *ψ* = 0.01, and host species richness of 3, 4, 5, 7 and 10 species. Table shows the standardized effect sizes of each metric. All parameter estimations were significant.

|  | **Scenario 1** |  |  |  | **Scenario 2** |  |  |
| --- | --- | --- | --- | --- | --- | --- | --- |
| Variable | Estimate effect on $R_0$ | $R^2$ | AIC | | Estimate effect on $R_0$ | $R^2$ | AIC |
| log(PD) | -0.44 | 0.89 | -3891 |  | -0.45 | 0.84 | -1206 |
| log(MPD) | -0.46 | 0.77 | 24 |  | -0.47 | 0.73 | 1452 |
| Ic | -0.05 | 0.22 | 6108 |  | -0.05 | 0.21 | 6765 |
| n | -0.13 | 0.40 | 4749 |  | -0.13 | 0.37 | 5647 |
| log(ED) | - | - | - |  | -0.47 | 0.70 | 1957 |
| Multivar (a) | 1.49-0.38log(MPD)-0.23n+0.04Ic | 0.91 | -4634 |  | 1.46-0.39log(MPD)-0.22n+0.03Ic | 0.85 | -1539 |
| Multivar (b) | - | - | - |  | 1.46-0.41log(ED)-0.30n+0.05Ic | 0.93 | -5428 |

Our full model was able to explain a high proportion of the variation (*R*^2^ = 0.91) in *R*_0_ across simulations, indicating that these relatively common phylogenetic indices are able to reliably capture the underlying disease transmission probabilities in our simulations.

#### Scenario 2: Single primary host

When we specify a primary host, the establishment probability of the disease (*c*) depends on the phylogenetic distance between the receiving and the primary host. Here, we quantify the phylogenetic isolation of the primary host using a metric of Evolutionary Distinctiveness (*ED*) with the fair proportions method (Isaac et al., 2007).

In the 3-taxon case (Figure 1), when the primary host is nested within the ingroup (Scenario 2a), the distance between the primary host (P) and the most proximate secondary host (S2), is the main driver of disease dynamics (Figures 1B and 1C), whereas the more distant secondary host (S1) contributes little to the outbreak potential because its further distance from the primary host reduces transmission probability. As the phylogenetic distance between the primary and proximate secondary host decreases (*PPd_P,S_*_2_ *→* 0), the model converges to a two-host system (P+S2 and S1). Similarly, when the primary host is sister to both secondary hosts (Scenario 2b), both secondary hosts are equidistant from the primary host (Figure 1D), and thus epidemiologically indistinguishable (here the system converges to a two-host system with P and S2+S1). However, as the distance between the primary host and secondary hosts (*PPd_P,S_*) increases, the system effectively converges on a one-host system, as successful transmission from the donor (P) to either secondary hosts becomes increasingly improbable.

In our single predictor regressions, effect sizes and model *R*^2^ are similar to those described for Scenario 1; however, *ED* is also strongly correlated with *R*_0_ (*R*^2^ = 0.70; Table 2), with *R*_0_ decreasing with increasing host *ED* (Figure 2E). Fitting the matching multiple regression model from Scenario 1 to data from Scenario 2, returns a notably worse fit (*R*^2^ = 0.85). However, when we substitute *MPD* for *ED*, we increased our explained variance to 93%, the highest across either scenario, and *ED* has the largest standardized effect size (Table 2). The model assuming FD transmission returned similar *R*^2^ (Appendix Section 6).

### 3.2 Option 2: Additive

Adding a species to a community does not effect the abundance of other species, such that the total number of individuals in the community increases.

In contrast to the models above assuming replacement, we here see a classic amplification effect (Figure 3) with species richness, *n*, positively correlated with *R*_0_. However, *MPD* and *ED* still show a diluting effect (Figure 3A, B and E), with absolute effect sizes comparable across all three variables in the multiple regression models. In the single predictor models, it is notable that the effect size of *MPD* is double that for *PD* in both scenarios, and *PD* alone explains little of the variation in *R*_0_ (Scenario 1: *R*^2^= 0.07 and, Scenario 2: *R*^2^=0.08). We suggest that the low explanatory power of *PD* is because it includes information on both *PPd* between species as well as species richness, which in this model option have an opposing relationship (classic amplification through *n*, phylogenetic dilution through *MPD*). The explanatory power of imbalance is larger than the estimated effect in option 1, and the sign of the relationship is switched—more balanced trees had a higher outbreak potential in a model of classic amplification. However, the contribution of tree balance to *R*_0_ is marginal in the multivariate model (Table 3).

**Table 3:**
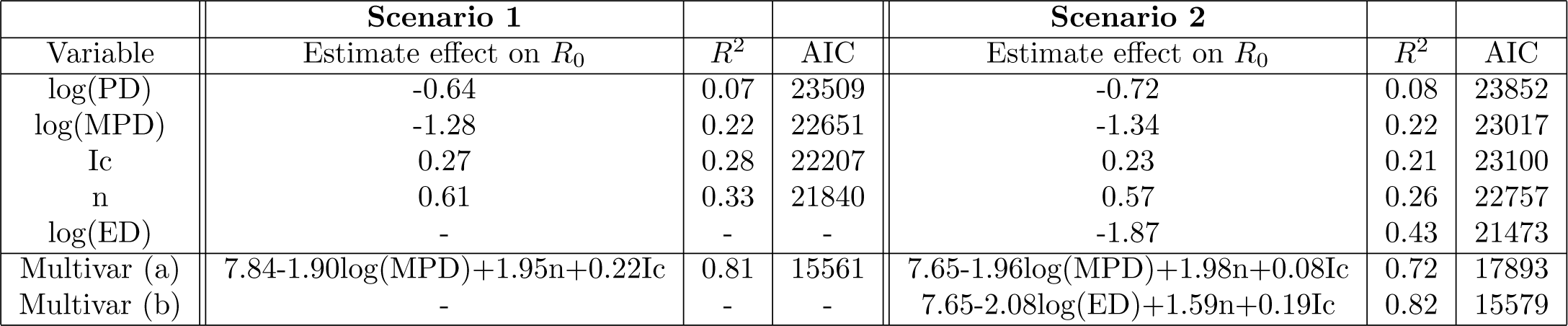
Models describing the variation in *R*_0_ with various phylogenetic metrics for simulations with model Option 2 (additive abundance) and *ν* = 0.1, *µ* = 0.2, *ψ* = 0.01, and host species richness of 3, 4, 5, 7 and 10 species. Table shows the standardized effects size of each metric. All parameter estimations were significant.

|  | **Scenario 1** |  |  |  | **Scenario 2** |  |  |
| --- | --- | --- | --- | --- | --- | --- | --- |
| Variable | Estimate effect on $R_0$ | $R^2$ | AIC | | Estimate effect on $R_0$ | $R^2$ | AIC |
| log(PD) | -0.64 | 0.07 | 23509 |  | -0.72 | 0.08 | 23852 |
| log(MPD) | -1.28 | 0.22 | 22651 |  | -1.34 | 0.22 | 23017 |
| Ic | 0.27 | 0.28 | 22207 |  | 0.23 | 0.21 | 23100 |
| n | 0.61 | 0.33 | 21840 |  | 0.57 | 0.26 | 22757 |
| log(ED) | - | - | - |  | -1.87 | 0.43 | 21473 |
| Multivar (a) | 7.84-1.90log(MPD)+1.95n+0.22Ic | 0.81 | 15561 |  | 7.65-1.96log(MPD)+1.98n+0.08Ic | 0.72 | 17893 |
| Multivar (b) | - | - | - |  | 7.65-2.08log(ED)+1.59n+0.19Ic | 0.82 | 15579 |

In scenario 2, we again find that the multivariate model with *ED* is favoured over that with *MPD* (*R*^2^=0.72 and 0.82 for the models with *MPD* and *ED*, respectively). Results for simulations assuming FD dynamics are the same for both additive and substitutive options (see Appendix section 6). Overall explanatory power for models with additive abundance (option 2) was lower than for models with substitutive abundance (option 1). Regression results are summarized in Table 3.

### 3.3 Application to Viruses and Bacteria of plants

We used the phylogenetic branch length transformations from Gilbert *et al*. (2012) for bacteria and viruses to weight probability of transmission success. We explored Scenario 1 only, as we did not have information on primary hosts. Assuming these empirically derived relationships, we show that host phylogeny is less important in determining community *R*_0_ for viruses (*R*^2^ = 0.49 for viruses vs *R*^2^ = 0.87 for bacteria; Table 4), which demonstrate a threshold-like relationship between *MPD* and *R*_0_ (Figure 4). Nonetheless, the multivariate models generally had high explanatory power across pathogen types (results for other pathogens are shown in the Appendix Table 2), with *MPD* the most important predictor variable.

**Table 4:**
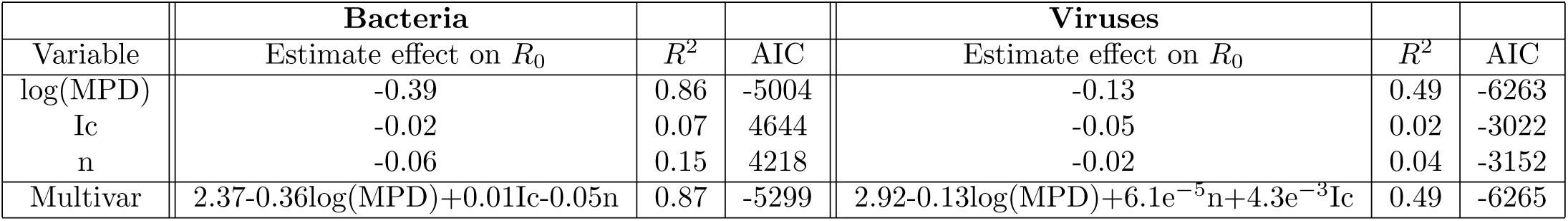
Regression coefficients as in Table 2, but assuming the phylogenetic transformation from Gilbert *et al*. (2012) for bacteria and viruses. Results are reported for Scenario 1 only, and are for simulations with *ν* = 0.1, *µ* = 0.2 and *ψ* = 0.01 and host species richness of 3, 4, 5, 7 and 10 species. The first three rows show estimates from the univariate regressions.

## 4 Discussion

We are witnessing a major rearrangement of biological diversity. Many species are threatened with extinction, while others are expanding their global distributions, sometimes invasively (IPBES, 2020; Barrett et al., 2018; IUCN, 2021). There is an urgent need to understand how these biotic changes might restructure host communities through altering environmental carrying capacities and species range-overlaps, and the cascading consequences of these changes on the maintenance and potential spillover of pathogens to novel hosts and to altered host communities (Daszak et al., 2001; Webb et al., 2002; Williams and Jackson, 2007; Faust et al., 2018). For example, host range expansions due to climate change may increase risk of pathogen sharing and spillover (Morales-Castilla et al., 2021). In addition, many host communities are also likely changing in diversity, with both decreases and increases in species richness common (Dornelas et al., 2014); yet the effects of biodiversity on disease prevalence remain controversial (Ostfeld and Keesing, 2012; Halliday and Rohr, 2019). Empirical studies provide evidence for both amplification and dilution effects, for example, increasing amphibian species richness has been linked to greater fungal pathogen prevalence (Becker and Zamudio, 2011), but reduced flatworm (*Ribeiroia*) prevalence (Johnson et al., 2012), and the mechanisms underlying these contrasting dynamics are often unclear.

Part of the explanation for the mixed support for dilution versus amplification effects likely reflects differences in disease competence among host species. The addition of highly competent hosts to the community will generally promote amplification, while the addition of lower-competent hosts or strong interspecific competitors that reduces the abundance of highly-competent hosts will tend to promote dilution (Cortez and Duffy, 2021). Thus, generating accurate predictions of how changes in host diversity will impact disease prevalence requires data on host competence, which we lack for most species. However, host phylogeny might provide a useful proxy (Gilbert and Parker, 2016; Gougherty and Davies, 2021).

Here, we present a novel framework that builds upon recent observations that disease transmission in multi-host systems is often strongly phylogenetically structured. Using simulations of a compartmental *SI* disease model with the assumption that the probability of successful transmission is inversely related to the pairwise phylogenetic distance between species, we show that simply by altering the phylogenetic relationships among hosts, a community’s disease outbreak potential (*R*_0_) can switch between amplifying (*R*_0_ *>* 1) to diluting (*R*_0_ *<* 1) pathogen invasion risk, while species richness remains unchanged. In our simulations, adding a closely related host to a community will more likely amplify a disease, while adding a distant relative will more likely dilute it. When we include primary-secondary host dynamics, we additionally show that it is not only the phylogenetic structure of the host community that determines the outbreak potential of the pathogen, but also the phylogenetic placement of the primary host. We were able to explain up to 93% of the variation in simulated community *R*_0_ using commonly-used phylogenetic diversity metrics that captured axes of tree topology and phylogenetic clustering, suggesting that these dimensions of host community structure might help in predicting disease spread. While the contribution of host phylogenetic structure to ecosystem functioning has been well studied (e.g. Cadotte et al., 2008) and there is increasing evidence for phylogenetic signal in disease transmission (Parker et al., 2015), these relationships have largely been explored independently; here, we provide a mechanistic model that unites them.

In our model, we make the well-supported assumption that the transmission of diseases among host species co-varies with the phylogenetic distance that separates them (Davies and Pedersen, 2008; Parker et al., 2015; Farrell and Davies, 2019b; Stephens et al., 2019; Wang et al., 2019; Gougherty and Davies, 2021), likely reflecting the tendency for closely related species to share similar immune systems, inherited, with modification, from a common ancestor. By extension, we assume the likelihood of establishment success of a disease in a novel host is correlated with the immuno-compatibility and thus phylogenetic identity of (donor and recipient) hosts (Davies and Pedersen, 2008; Kuiken et al., 2006; Streicker et al., 2010; Gilbert et al., 2012; Huang et al., 2014). It has already been shown that the genetic diversity of host populations can affect disease outbreak potential (Altizer et al., 2003) and that *R*_0_ may be inversely related to the number of host genotypes (Lively, 2010). We extend this reasoning to encompass host phylogenetic diversity and show that under assumptions of phylogenetic conservatism in host competence, community *R*_0_ is lower in simulated communities with greater host phylogenetic diversity, indexed by the phylogenetic pairwise distances separating hosts in the community.

We present a model that allows for a *phylogenetic dilution* effect, by assuming an inverse relationship between phylogenetic distance between hosts and risk of pathogen sharing. In our model, we include a phylogenetic scaling parameter, *α*, that allows us to weight the relative importance of phylogenetic distance. We suggest *α* should be considered as a free parameter, best estimated from the data, and we discuss an example below using a scaling transformation from Gilbert et al. (2012). Importantly, however, any shape can be fit, depending on the host-pathogen system (see Appendix Section 5). It would be straightforward to transform branch lengths to match to alternative evolutionary models, such as the Ornstein–Uhlenbeck (O-U) process (Martins and Hansen, 1997), or more complex, rate variable models.

We explored dynamics across two alternative scenarios. First, we assume transmission success is simply weighted by the phylogenetic distance between donor and recipient hosts (Scenario 1), which we suggest might resemble disease dynamics in pathogen systems such as Rabies virus, characterised by frequent host jumps and fast mutation rates, allowing it to co-evolve with its host (Woolhouse et al., 2001; Mollentze et al., 2014). However, the majority of disease systems likely have more complex primary-secondary host dynamics, in which one or a few host species are the primary (or reservoir) hosts, and host competence co-varies with phylogenetic distance between the secondary (recipient) and primary host (Scenario 2). We highlight bovine tuberculosis (bTB) in buffalo as one exemplar disease system that fits this latter scenario. In Africa, bTB, which originated from cattle imported from Europe, is now endemic in African buffalo, but frequently spills over to other mammal hosts (Michel et al., 2006).

In both modelled scenarios with substitutive abundance (option 1), an increase in species richness (*n*) decreases *R*_0_. The inclusion of a low competent host in a population with a constant contact rate per time unit (frequency dependent transmission) can lead to a reduction in contacts among high competent hosts, resulting in a lower overall disease prevalence (Ostfeld and Keesing, 2000; Dobson, 2004; Rudolf and Antonovics, 2005; Keesing et al., 2006). Disease dilution is also possible with density dependent transmission if we assume the total number of individuals in the community remains constant (i.e. substitutive abundance), such that added hosts replace contacts among more competent hosts. Phylogenetic tree shape was weakly correlated with disease outbreak potential in both scenarios: more balanced trees reduce community *R*_0_. A highly imbalanced tree is characterized by one or a few clades of very closely related species, which are more likely to have similar immune-competence, and thus easily share a disease, we suggest this parallels the *phylogenetic clade effect* discussed by Wang et al. (2019). However, in the absence of an assigned primary host, the best predictor of community *R*_0_ (in combination with *S* and *Ic*) was the mean of the pairwise phylogenetic distance between hosts in the community (*MPD*). Communities composed of more closely related hosts had higher outbreak potentials than communities composed of more distantly related hosts because the average transmission success between recipient and donor hosts will be relatively high in the former and low in the latter. When we include a primary host in the model (Scenario 2), we are again able to explain most of the variation in community *R*_0_ using just species richness, tree imbalance, and *MPD*. However, the model substituting *MPD* for the evolutionary isolation (measured as Evolutionary Distinctiveness; *ED*) of the primary host was a better fit to the data. In this modelled scenario, community *R*_0_ is lower when the primary host is phylogenetically isolated from secondary hosts, and primary host ED is more important in mediating disease dynamics than overall host phylogenetic community structure.

Notably, when we assume an increase in species richness is additive with respect to community abundance (option 2), the effect of species richness (*n*) on *R*_0_ switched from diluting to amplifying, while the phylogenetic dilution effect of *MPD* is largely unchanged. This modelled scenario illustrates how both amplification and dilution effects can operate simultaneously on different axes of community structure. Additionally, we observe a drop in explanatory power of ED in scenario 2, suggesting that when encounters are additive, and thus amplifying, the location of the primary host is of less importance in mediating outbreak potential.

The shape of the relationship between host phylogeny and disease outbreak potential, as well as the relative importance of the different axes of phylogenetic structure we explore, reflect both the underlying structure of our model and the parameter values we fit to it. To examine how these relationships might change under alternative functions describing the phylogenetic signal in the probability of pest sharing, we also fit our model using the functions from Gilbert et al. (2012) estimated from empirical data for viruses and bacteria on plants. As we do not have information on primary hosts, we fit a model for scenario 1 only. For both pathogen types, we can see the relationship between *R*_0_ and *MPD* is qualitatively different from that shown in our simulations assuming a simpler phylogenetic scaling. Perhaps most notably, the diluting effect of *MPD* is only apparent at much larger values, reflecting the ease with which many generalist pathogens can jump between closely related species (Schatz and Park, 2021). Phylogenetic constraints to host shifts are only apparent at high *MPD*, and for viruses there is evidence of a threshold below which host jumps are effectively unconstrained by phylogeny. Viruses are recognised as being able to make relatively large host jumps (Woolhouse et al., 2005; Park et al., 2018), therefore it may not be so surprising that we also find in our model that host community phylogenetic structure is less important in mediating their outbreak potential.

We recognise that our model, like all models, is an over-simplification, and the high *R*^2^ from our regressions reflect some of the many simplifying assumptions we impose, including equal host population sizes and symmetries in contact rates. Importantly, we also modelled the intraspecific and interspecific contact rates as equal, which would have had an influence on the relative magnitude of amplification/dilution effects (Dobson, 2004; Joseph et al., 2013); however, this is easily modified if different assumptions on host contacts were a better fit to our understanding of a particular empirical system. In Appendix section 4 we show how *R*^2^ from regressions increases when we switch to more realistic contact structures where the interspecific contact rates are reduced relative to intraspecific contacts. Our models are also deterministic, and thus ignore potential for stochastic dynamics, which can have significant impacts on disease outbreaks dynamics (Wolfe et al., 2005). We thus see many exciting extensions to the model framework we present here.

Our simulations, based upon the reasonable assumptions that spillover of pathogens is more likely between closely related hosts, highlight how the phylogenetic structure of host communities might additionally mediate risk of disease outbreaks: communities that are more phylogenetically diverse are also more likely to reduce disease outbreak potential (see also Davies and Pedersen, 2008; Parker et al., 2015; Farrell and Davies, 2019b; Wang et al., 2019, 2021). However, there is evidence that more evolutionarily distinct species—those adding the greatest phylogenetic diversity to communities—might also be more at risk of extinction (Daru et al., 2013), suggesting that extinction might shift communities towards increased potential for disease outbreaks. In addition, the non-random distribution of extinction on the tree-of-life suggests that, at least at local scales where disease dynamics play out, we risk losing a disproportionate amount of phylogenetic diversity if currently threatened species are lost to extinction (Huang et al., 2012). Thus, we may be most at risk of losing the species and community attributes most likely to suppress disease, while the species less at risk of extinction tend to be those associated with faster life-histories and higher disease competence due to lower investment in pathogen defense (Joseph et al., 2013; Gibb et al., 2020; Halliday et al., 2020; Valenzuela-Sánchez et al., 2021).

## Acknowledgements

We would like to thank S. Otto for reviewing an earlier version of this manuscript and her advice on modelling approaches. We also thank Ailene MacPherson for her help with the modelling. This research was supported by an NSERC Discovery grant awarded to TJD.

## Author contributions

Manuscripts was conceptualized by T.J. Davies and M.E.M. Toorians. MEMT designed and analysed the model with feedback from I.M. Smallegange and TJD. MEMT wrote the paper with TJD. IMS contributed to manuscript edits.

## Data accessibility

Data has been generated in simulations in R scripts that have been deposited on https://github.com/Marjoleintoorians/PhyloSIModel

