## Supplementary text for "Host community structure can shape pathogen outbreak dynamics through a phylogenetic dilution effect"

March 4, 2024

#### 1 Derivation of the multi-host model

For a system with $n$ hosts:

$$\frac{dS_i}{dt} = b_i S_i - \sum_{j=1}^{n} \beta_{ij} I_j p_{ij} - \nu_i S_i \tag{1a}$$

$$\frac{dI_i}{dt} = \sum_{j=1}^{n} \beta_{ij} I_j p_{ij} - (\nu_i + \mu_i) I_i \tag{1b}$$

Where $p_{ij} = \frac{S_i}{N_j}$ for Density-dependent transmission and $p_{ij} = \frac{S_i}{N}$ for Frequency-Dependent transmission. We will here show the derivation of the Next Generation Matrix (NGM) for the calculation of $R_0$ for both density- and frequency dependence. First, we establish the difference between the two transmission modes through the contact rates.

#### 1.1 Density-dependence

Here, the rate of transmission is determined by the density of hosts. Per capita transmission rate is:

$$\beta_{ij}(N_j) = \kappa_{ij}(N_j) \cdot c_{ij} \tag{2}$$

The contact rate ($\kappa_{ij}$) is a density dependent rate per $i$ individual, therefore the Eqn is:

$$\kappa_{ij}(N_j) = \xi_i N_j \tag{3}$$

Here, we show the derivation of the NGM of main text Eqn 3 for a simple 2-host model:

$$\frac{dS_1}{dt} = -\beta_{11} S_1 I_1 - \beta_{12} S_1 I_2 - \nu_1 S_1 \tag{4a}$$

$$\frac{dS_2}{dt} = -\beta_{21} S_2 I_1 - \beta_{22} S_2 I_2 - \nu_2 S_2 \tag{4b}$$

$$\frac{dI_1}{dt} = \beta_{11} S_1 I_1 + \beta_{12} S_1 I_2 - (\nu_1 + \mu_1) I_1 \tag{4c}$$

$$\frac{dI_2}{dt} = \beta_{21} S_2 I_1 + \beta_{22} S_2 I_2 - (\nu_2 + \mu_2) I_2 \tag{4d}$$

Now, to calculate the $R_0$ we linearize around the disease-free equilibrium and find the invasion criterion, which is the equivalent of the $R_{0,total}$ (Diekmann et al., 2010; Hurford et al., 2010). We begin by calculating the Jacobian matrix:

$$J = \begin{bmatrix} \frac{\partial \dot{I_1}}{\partial I_1} & \frac{\partial \dot{I_1}}{\partial I_2} \\ \frac{\partial \dot{I_2}}{\partial I_1} & \frac{\partial \dot{I_2}}{\partial I_2} \end{bmatrix} = \begin{bmatrix} \beta_{11} S_1 - (\mu_1 + \nu_1) & \beta_{12} S_1 \\ \beta_{21} S_2 & \beta_{22} S_2 - (\mu_2 + \nu_2) \end{bmatrix} \tag{5}$$

Then, we separate the Jacobian matrix into two components: The transition matrix ($\Sigma$) and the Transmission matrix (**T**):

$$T = \begin{bmatrix} \beta_{11} S_1 & \beta_{12} S_1 \\ \beta_{21} S_2 & \beta_{22} S_2 \end{bmatrix} \tag{6}$$

and

$$\Sigma = \begin{bmatrix} -(\mu_1 + \nu_1) & 0 \\ 0 & -(\mu_2 + \nu_2) \end{bmatrix} \tag{7}$$

Then, we calculate the Next-Generation-Matrix **K** as:

$$K = -T\Sigma^{-1} \begin{bmatrix} \frac{\beta_{11}S_1}{\mu_1+\nu_1} & \frac{\beta_{12}S_1}{\mu_2+\nu_2} \\ \frac{\beta_{21}S_2}{\mu_1+\nu_1} & \frac{\beta_{22}S_2}{\mu_2+\nu_2} \end{bmatrix} \tag{8}$$

The maximum eigenvalue of this matrix is the $R_{0,total}$ of this community of hosts.

#### 1.2 Frequency-dependence

For frequency dependent transmission (sexually transmitted diseases or approximations of vector-transmitted diseases), the density of host does not determine the transmission. Transmission is a constant rate per time unit.

$$\beta_{ij} = \kappa_i \cdot c_{ij} \tag{9}$$

The contact rate ($\kappa_{ij}$) is a density dependent rate per $i$ individual, therefore the Eqn is:

$$\kappa_i = \xi_i \tag{10}$$

Here, we show the derivation of the NGM similar to above, but using frequency-dependent transmission in a 2-host model:

$$\frac{dS_1}{dt} = -\beta_{11}S_1\frac{I_1}{N} - \beta_{12}S_1\frac{I_2}{N} - \nu_1 S_1 \tag{11a}$$

$$\frac{dS_2}{dt} = -\beta_{21}S_2\frac{I_1}{N} - \beta_{22}S_2\frac{I_2}{N} - \nu_2 S_2 \tag{11b}$$

$$\frac{dI_1}{dt} = \beta_{11}S_1\frac{I_1}{N} + \beta_{12}S_1\frac{I_2}{N} - (\nu_1 + \mu_1)I_1 \tag{11c}$$

$$\frac{dI_2}{dt} = \beta_{21}S_2\frac{I_1}{N} + \beta_{22}S_2\frac{I_2}{N} - (\nu_2 + \mu_2)I_2 \tag{11d}$$

$$\tag{11e}$$

From this set of equations, we calculate the Jacobian matrix:

$$J = \begin{bmatrix} \frac{\partial \dot{I_1}}{\partial I_1} & \frac{\partial \dot{I_1}}{\partial I_2} \\ \frac{\partial \dot{I_2}}{\partial I_1} & \frac{\partial \dot{I_2}}{\partial I_2} \end{bmatrix} = \begin{bmatrix} \beta_{11}\frac{S_1}{N} - (\mu_1 + \nu_1) & \beta_{12}\frac{S_1}{N} \\ \beta_{21}\frac{S_2}{N} & \beta_{22}\frac{S_2}{N} - (\mu_2 + \nu_2) \end{bmatrix} \tag{12}$$

We can then again separate the matrix into a transmission matrix (T) and a transition matrix ($\Sigma$):

$$T = \begin{bmatrix} \beta_{11}\frac{S_1}{N} & \beta_{12}\frac{S_1}{N} \\ \beta_{21}\frac{S_2}{N} & \beta_{22}\frac{S_2}{N} \end{bmatrix} \tag{13}$$

and

$$\Sigma = \begin{bmatrix} -(\mu_1 + \nu_1) & 0 \\ 0 & -(\mu_2 + \nu_2) \end{bmatrix} \tag{14}$$

Then, we calculate the Next-Generation-Matrix $\mathbf{K}$ as:

$$K = -T\Sigma^{-1} = \begin{bmatrix} \frac{\beta_{11}\frac{S_1}{N}}{\nu_1+\mu_1} & \frac{\beta_{12}\frac{S_1}{N}}{\nu_2+\mu_2} \\ \frac{\beta_{21}\frac{S_2}{N}}{\nu_1+\mu_1} & \frac{\beta_{22}\frac{S_2}{N}}{\nu_2+\mu_2} \end{bmatrix} \tag{15}$$

Where the maximum eigenvalue of matrix $\mathbf{K}$ is the $R_0$ of the community of hosts.

### 2 Phylo-SI model validation

#### 2.1 Community simulations

Host communities were simulated using a pure-birth Yule-process with the *pbtree()* function from the *Phytools* library in R (v. 4.3.2). This function simulated trees with varying tree depth and an assigned number of species ($n$). The birth rate ($b$) determines branching times (Figure 1), and thus the phylogenetic diversity ($PD$) of the tree comprising the simulated community; increasing $b$ reduces $PD$ (Figure 2A) and $MPD$ (Figure 2B).This coincides with an increase in $R_0$ (Figure 2C).

### 3 PCA

#### 3.1 Option 1

Using simulations, we show that $R_0$ varies with both the richness of hosts and the topology of the phylogenetic tree that connects them (Figure 3, left panel). In a Principal Component Analysis (PCA) for models with option 1, we show that 91% of variation falls along two independent axes: PCA1 - 66.85%- with weights predominantly reflecting MPD and PD, and PCA2 - 23.77%- which captures variation in tree imbalance (Figure 3, left panel). The PCA for

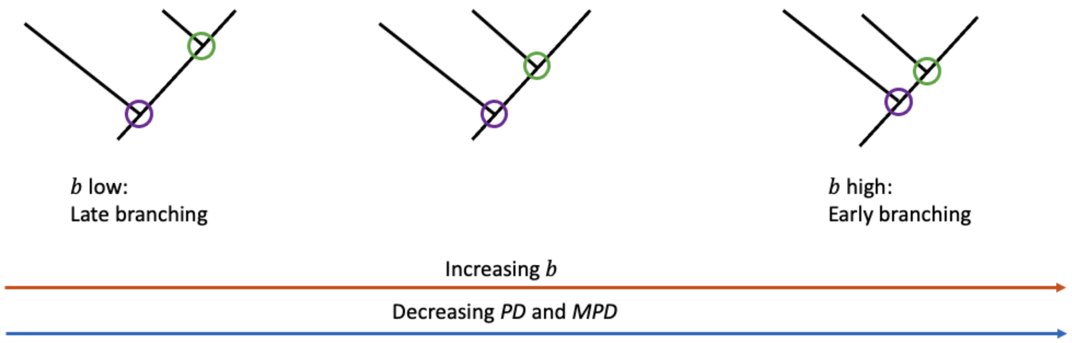


Figure 1: Illustration showing effect of varying birth rate, $b$, for a three-species tree. Increasing $b$ decreases branching time between speciation events, therefore the species (tips of the tree) are more closely related, when we allow for the tree depth to vary and holding number of species constant. The purple circle represents the first speciation event, the green circle the second speciation event in this 3-species tree.

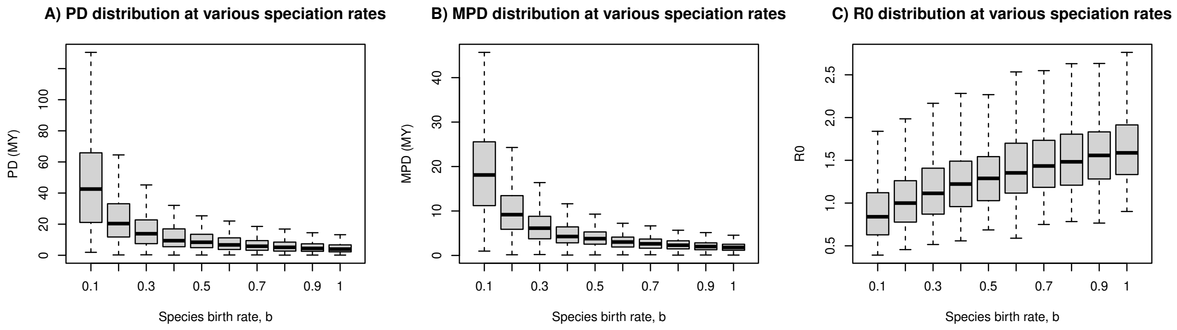


Figure 2: Boxplots showing host tree phylogenetic diversity (A), Mean Pairwise phylogenetic Distance (B) and corresponding community $R_0$ (C), for communities of $n$ = 3,4,5,7 and 10 species, simulated assuming different birth rates, $b$. Data were generated from simulations using $\psi = 0.01$, $\nu = 0.1$, $\mu = 0.2$. 100 communities were simulated per species birth rate value, $b$. Graphs look the same for Options 1 and 2.

scenario 2 with multiple hosts (where the primary host is randomly assigned in each simulation) reveals that 89 % of the variation is distributed over the first two axes (Figure 3, right panel). $ED$ is closely aligned with $PD$ and $MPD$.

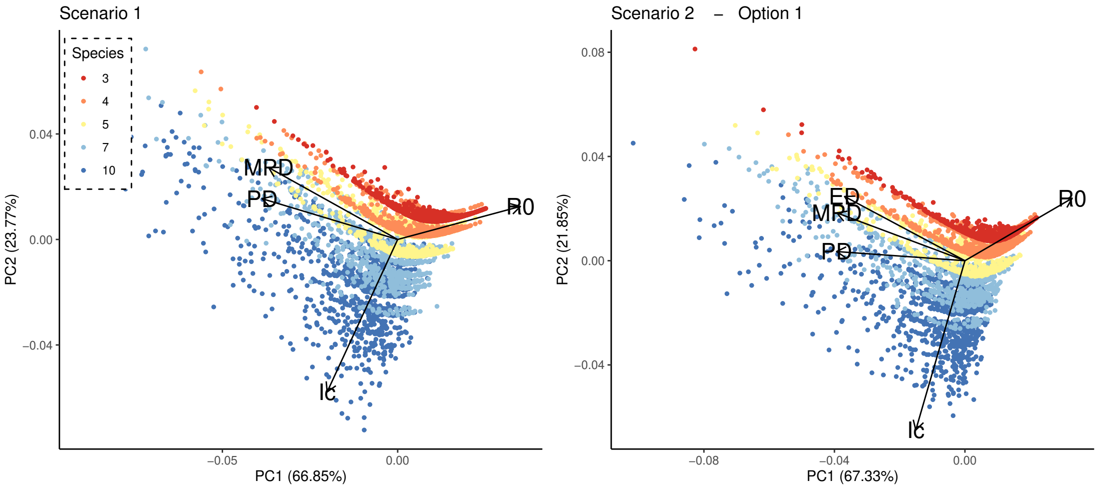


Figure 3: Principal Component Analysis for DD model using Option 1 for Scenarios 1 (left) and 2 (right) with $\nu = 0.1$, $\mu = 0.2$ and $\psi = 0.01$. Colours indicate simulated communities with different host richness ($n$=3,4,5,7,10).

#### 3.2 Option 2

Repeating this in a model with additive abundance, we show that 93% of the variation in Scenario 1 falls over two axes, 54.66% reflecting PD and MPD, and 38.04% tree topology, imbalance (Figure 4). In the second scenario, the total variation can be explained by these two axes (61.1% for the axes reflecting PD,MPD and ED, and 29.2% for the tree topology) for a total of 90%, showing an increase in this scenario including a primary reservoir, opposite to what we have seen in models with option 1. The greatest difference between options 1 and 2 is the change in direction of Ic. However, the contribution is only small (see Tables 3 and 4 in main text). We see a clear difference in the distribution of species richness, which is due to the dilutings versus amplifyingg behaviour of the model.

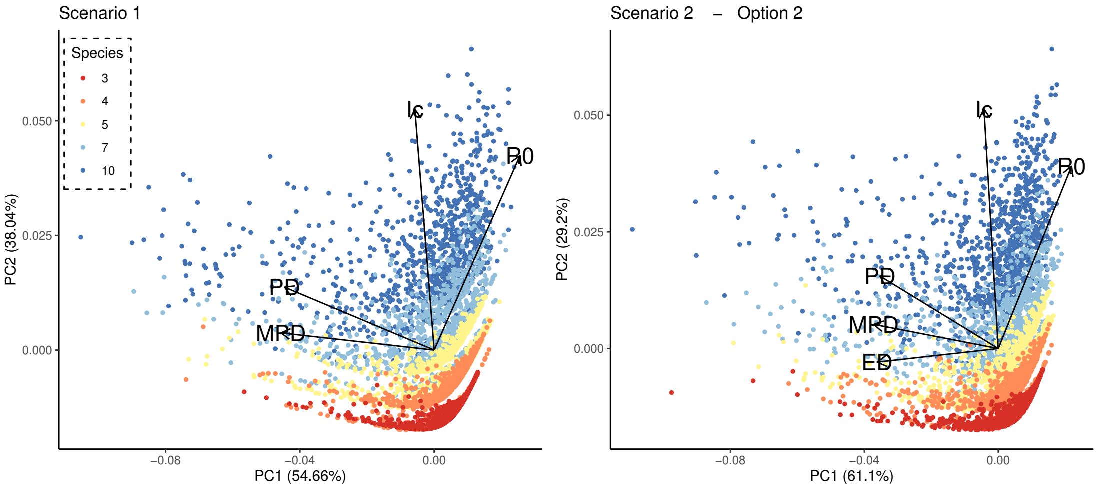


Figure 4: Principal Component Analysis for DD model using Option 2 for Scenarios 1 (left) and 2 (right) with $\nu = 0.1$, $\mu = 0.2$ and $\psi = 0.01$. Colours indicate simulated communities with different host richness ($n$=3,4,5,7,10).

### 4 Alternative contact rate structures

To explore more realistic contact rates, we generated an alternative set of simulations with interspecific contact rates lower than intraspecific contact rates. Here, we use *Kronecker's* $\delta$ function for the contact rate:

$$\kappa(N) = \kappa_0(\delta_{ij} + (1 - \nu_{ij})\alpha) \cdot f(N) \tag{16}$$

when $i = j$, then $\nu = 1$, if $i \neq j$, $\nu = 0$. $\kappa_0$ represents the intraspecific contacts, and $\alpha$ represents the interspecific contacts (note that here $\nu$ is Kronecker's $\nu$, not the natural mortality rate used in the $SI$ model in the main text). The function $f(N)$ determines the nature of the transmission. In density-dependent systems, $f(N) = 1$, whereas in frequency-dependent systems, such as used in the model in the main text, $f(N) = \frac{1}{N}$. Here, we explore two other contact structures for both scenarios:

#### 4.1 Interspecific contact rate = 0.1 x Intraspecific contact rate

In this model, we assume that the interspecific contacts are 10 times less likely than intraspecific contacts. Here, the interspecific contact rates are set to 0.1, and the intraspecific contact rates are set to 1.

##### 4.1.1 DD model option 1

Model dynamics are similar to the main text Figure 2. These results are shown in Figure 6. Multivariate model for Scenario 1 is $R_0 = 1.34 - 0.34log(MPD) + 0.03Ic - 0.21n$, with an $R^2 = 0.91$. For Scenario 2, model (a) is $R_0 = 1.31 - 0.35log(MPD) + 0.02Ic - 0.20n$ with $R^2 = 0.85$, and model (b) $R_0 = 1.31 - 0.36log(ED) + 0.04Ic - 0.27n$ with $R^2 = 0.93$.

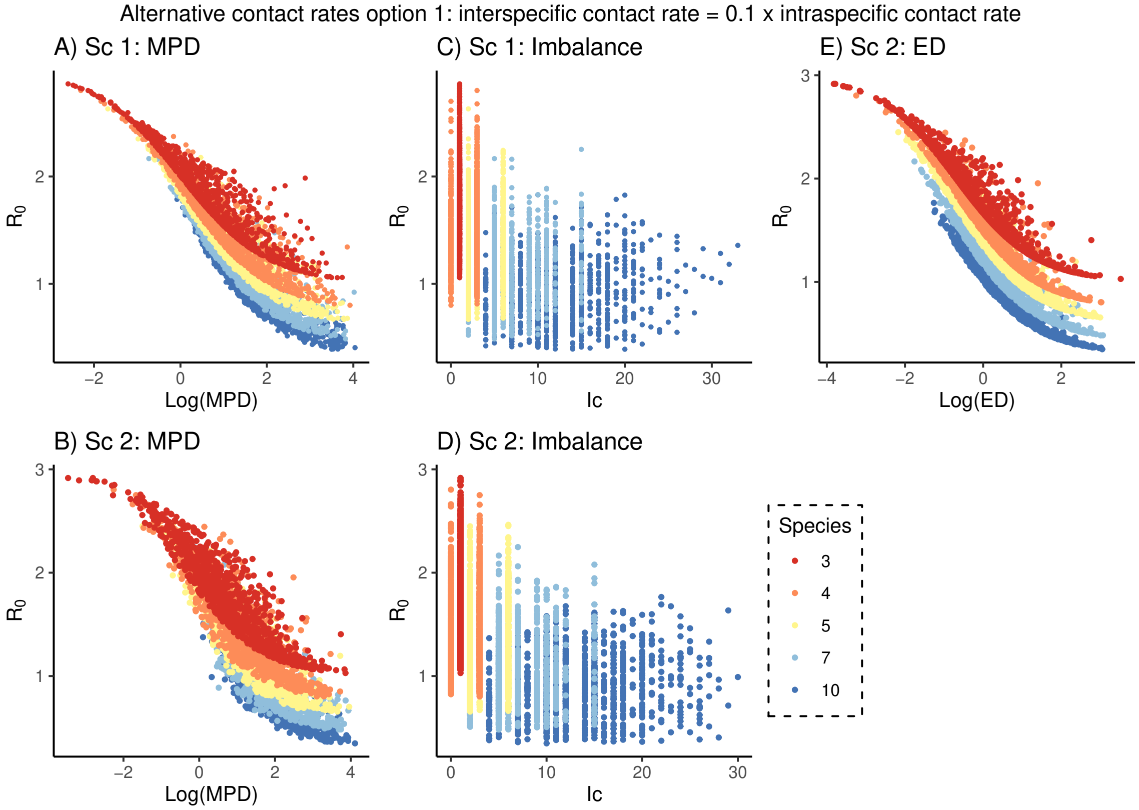


Figure 5: Model Option 1 with alternative contact rates. A,B: Mean Pairwise Distance ($MPD$) of the community is inversely related to $R_0$ in Scenario 1 (A) and Scenario 2 (B). C,D: Phylogenetic Imbalance of the tree connecting the host species has variable effects on $R_0$. E: Evolutionary Distinctiveness of primary host in (Scenario 2) decreases $R_0$. Data from simulations with $\psi = 0.01$, $\nu = 0.1$, $\mu = 0.2$. Colours indicate simulated communities with different host richness ($n$=3,4,5,7,10). Interspecific contact rates, $\kappa_{ij}$, are 0.1, while intraspecific contact rates, $\kappa_{ii}$, are 1.

###### 4.1.2 DD model option 2

Results are very similar to main text Figure 3. Now, $R_0$ no longer drops below 1, thus, the pathogen always causes an epidemic. The pathogen predominantly spreads within the primary reservoir, and there is less spillover. This is shown in Figure 6. Multivariate model for Scenario 1 is $R_0 = 7.06 - 1.71 log(MPD) + 0.20 Ic - 1.76n$, with an $R^2 = 0.81$. For Scenario 2, model (a) is $R_0 = 6.88 - 1.77 log(MPD) + 0.07 Ic + 1.78n$ with $R^2 = 0.72$, and model (b) $R_0 = 6.88 - 1.87 log(ED) + 0.17 Ic - 1.43n$ with $R^2 = 0.82$. In these models there is a larger effect of species richness, $n$, on $R_0$ (Figure 6, in colours), with an increase in species richness dramatically reducing $R_0$ relative to the model assuming equal contact rates presented in the main text. Because interspecific transmission now contributes ten times less to total disease transmission, adding a novel species to the community adds only a small amount of interspecific transmission to the community. We only show results for model option 1.

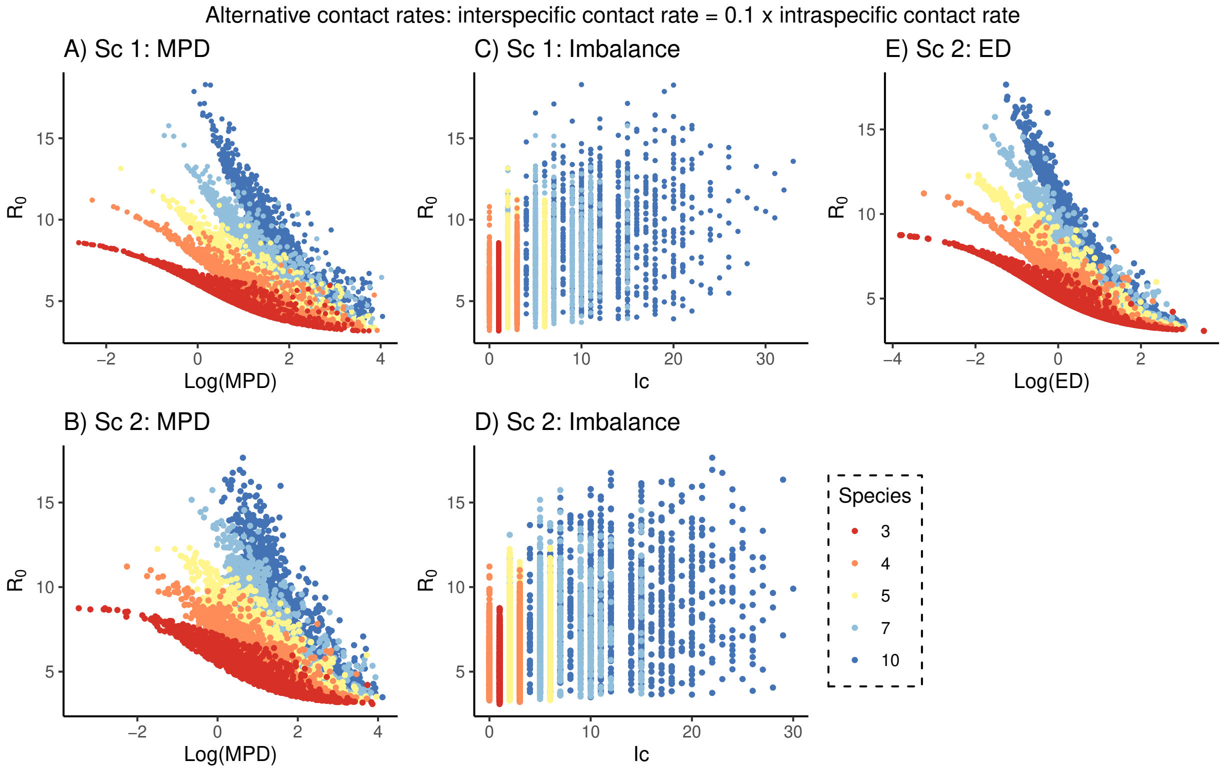


Figure 6: Model Option 2 with alternative contact rates. A,B: Mean Pairwise Distance ($MPD$) of the community is inversely related to $R_0$ in Scenario 1 (A) and Scenario 2 (B). C,D: Phylogenetic Imbalance of the tree connecting the host species has variable effects on $R_0$. E: Evolutionary Distinctiveness of primary host in (Scenario 2) decreases $R_0$. Data from simulations with $\psi = 0.01$, $\nu = 0.1$, $\mu = 0.2$. Colours indicate simulated communities with different host richness ($n$=3,4,5,7,10). Interspecific contact rates, $\kappa_{ij}$, are 0.1, while intraspecific contact rates, $\kappa_{ii}$, are 1.

#### 4.2 Interspecific contact rates drawn from a uniform distribution

Here, interspecific contact rates, $\kappa_{ij}$, were drawn from $U[0.1, 0.5]$. This allows greater interspecific contact rates than in the previous simulations, reducing $R_0$, as shown in Figure 7. We only show results for model option 1. Multivariate model for Scenario 1 is $R_0 = 1.34 - 0.34log(MPD) + 0.03Ic - 0.21n$, with an $R^2 = 0.91$. For Scenario 2, model (a) is $R_0 = 1.31 - 0.35log(MPD) + 0.02Ic - 0.20n$ with $R^2 = 0.85$, and model (b) $R_0 = 1.31 - 0.36log(ED) + 0.04Ic - 0.27n$ with $R^2$=0.93. In these models there is a larger effect of $ED$ on $R_0$ (Figure 7E) relative to the model assuming equal contact rates presented in the main text. These models shows moderate effects of the interspecific transmission, and may be considered as an intermediate between the model presented in the main text and model above, assuming a 10-fold difference in contact rates between versus within host species.

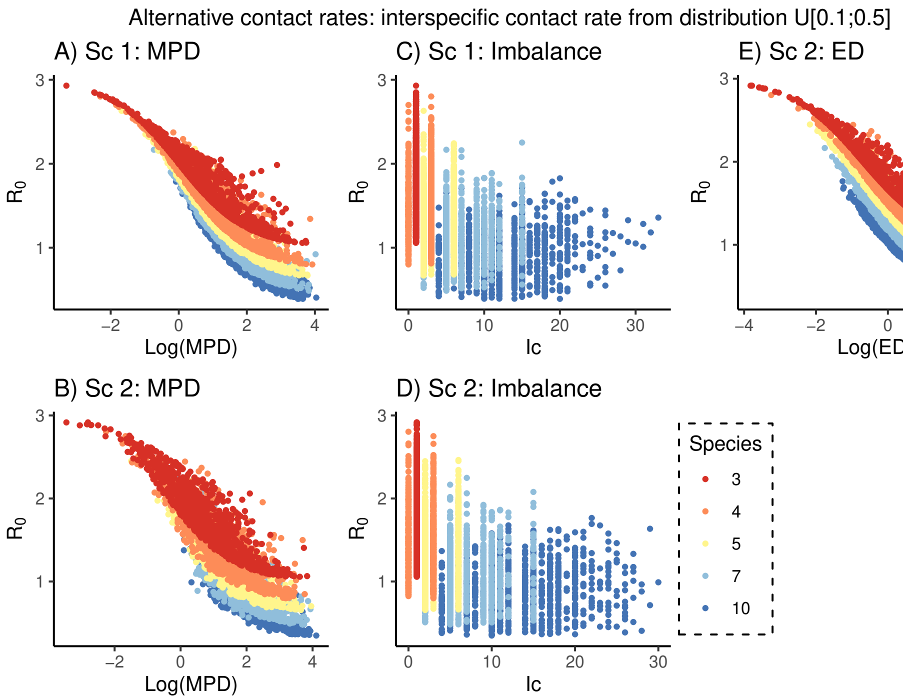


Figure 7: Model Option 1 with alternative contact rates. A,B: Mean Pairwise Distance ($MPD$) of the community is inversely related to $R_0$ in Scenario 1 (A) and Scenario 2 (B). C,D: Phylogenetic Imbalance of the tree connecting host species has variable effects on $R_0$. E: Evolutionary Distinctiveness of the primary host in Scenario 2 decreases $R_0$. Data from simulations with $\psi = 0.01$, $\nu = 0.1$, $\mu = 0.2$. Colours indicate simulated communities with different host richness ($n$=3,4,5,7,10). $\kappa_{ij}$, were drawn from a uniform distribution ranging from 0.1-0.5. Intraspecific contact rates, $\kappa_{ii}$, are 1.

### 5 Negative exponential probability of successful transmission

The model we present in the main text describes an inverse relationship between transmission potential and phylogenetic distance between hosts (Lively, 2010; Poullain and Nuismer, 2012); however, the form of this relationship may differ between host species and pathogen-types (Gilbert et al., 2012). Here, we consider an alternative functional form of this relationship based on empirical work by Gilbert et al. (2012), assuming an exponential decrease in successful transmission with phylogenetic distance, $PPd$. Regression coefficients matching to those presented in the main text are shown in table 1. We show results for only DD and option 1.

| | Scenario 1 | | Scenario 2 | |
|---|---|---|---|---|
| Variable | Estimate effect on $R_0$ | $R^2$ | Estimate effect on $R_0$ | $R^2$ |
| log(PD) | -0.38 | 0.75 | -0.40 | 0.71 |
| log(MPD) | -0.38 | 0.61 | -0.40 | 0.58 |
| Ic | -0.04 | 0.23 | -0.05 | 0.24 |
| n | -0.12 | 0.41 | -0.13 | 0.41 |
| log(ED) | - | - | -0.40 | 0.55 |
| Multivariate (a) | 1.08-0.30log(MPD)-0.24n+0.04Ic | 0.79 | 0.99-0.32log(MPD)+ 0.02Ic -0.24n | 0.75 |
| Multivariate (b) | - | - | 0.99-0.34log(ED)-0.31n+0.04Ic | 0.83 |

Table 1: Regressions coefficients assuming a negative exponential relationship between transmission potential and phylogenetic distance. Univariate and multivariate regressions, describing the variation in $R_0$ using various phylogenetic metrics for Scenario 1 and 2, using $\nu = 0.1$, $\mu = 0.2$ and $\psi = 0.01$ in simulated communities with a host species richness of 3, 4, 5, 7 and 10 species. Table shows the standardized effects size of each metric and model explanatory power for $R_0$. All parameter estimations were significant.

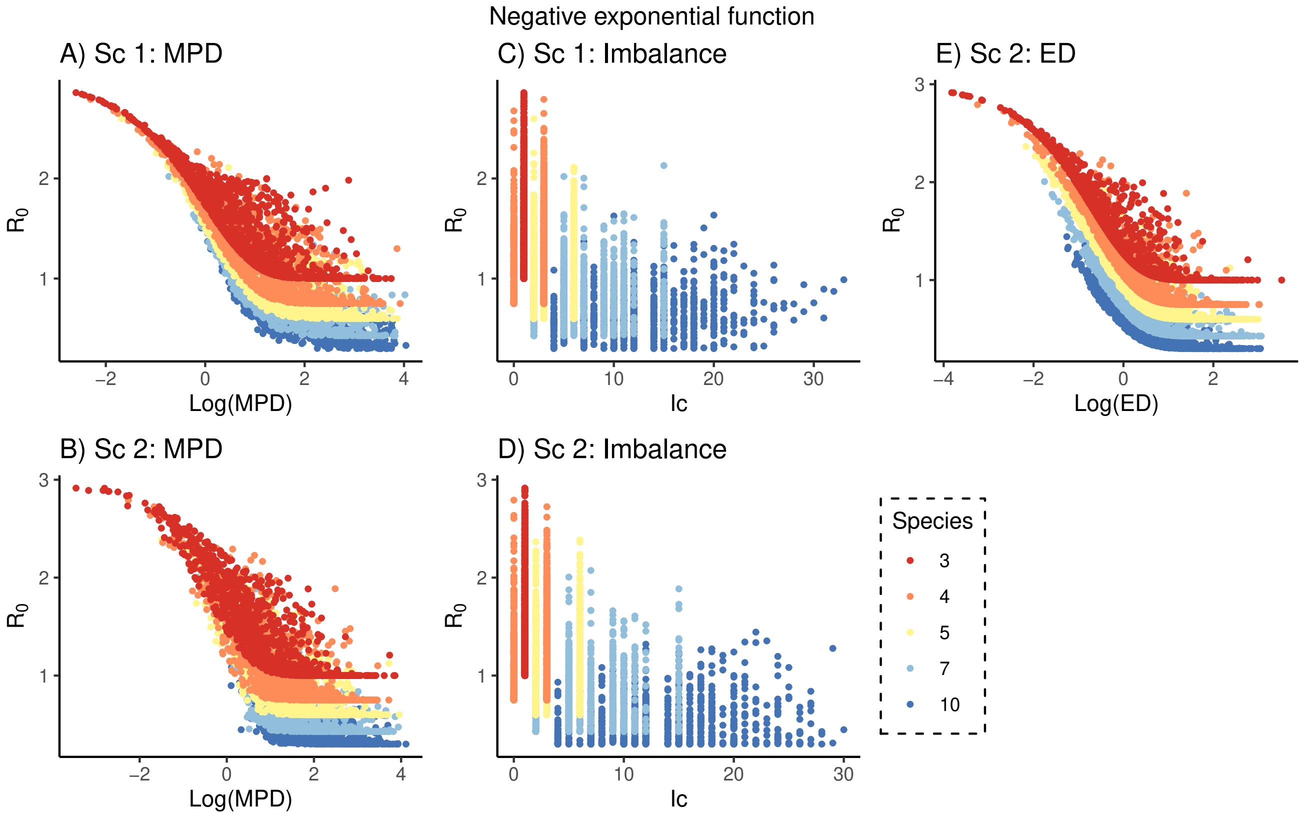


Figure 8: Negative exponential relationship between transmission, $c$, and $PPd$. DD model with option 1 (Replacing contacts). A,B: Mean Pairwise Distance ($MPD$) of the community is inversely related to $R_0$ in both scenarios. C,D: Phylogenetic Imbalance of the tree connecting the host species community has variable effects $R_0$. E: Evolutionary Distinctiveness of primary host in Scenario 2 decreases $R_0$. Data from simulations with $\psi = 0.01$, $\nu = 0.1$, $\mu = 0.2$. Colours indicate simulated communities with different host richness ($n$=3,4,5,7,10). Intra- and interspecific contact rates, $\kappa_{ij}$, are 1.

### 6 Frequency-dependent model

In the main text we focused on density-dependent models only. However, frequency-dependent (FD) models are commonly-used in disease-ecology for their simplicity to model pathogens whose transmission does not rely on host density. For example, sexually-transmitted or vector-transmitted diseases (De Jong et al., 1995; Keeling and Rohani, 2011).

The FD model is described in Section 1.2, and here we show a model option where we replace contacts. For FD, the contact structure (replacing or additive) does not have an effect on $R_0$, therefore option 1 and 2 yield the same results (See Figure 9). Here, $R_0$ values are a order of magnitude lower as we use the same parameter values, but modelling in FD. Per usual, transmission values need to be adjusted accordingly such that $R_0$ is the same for DD and FD at a constant population size, $N$ (De Jong et al., 1995). This model will always lead to dilution, as predicted by theoretical studies (Rudolf and Antonovics, 2005). Regressions show different coefficients for the variables' effect on $R_0$, but the $R^2$ relationship remains the same: Model A for Scenario 1 $R^2 = 0.91$. For Scenario 2, model A drops down to $R^2 = 0.85$, but in model B Scenario 2 goes back up to $R^2$=0.93, similarly showing the importance of the community structure and location of the primary host. These are the same values as in DD option 1, as both portray similar diluting effects (due to replacing the contacts when increasing species richness).

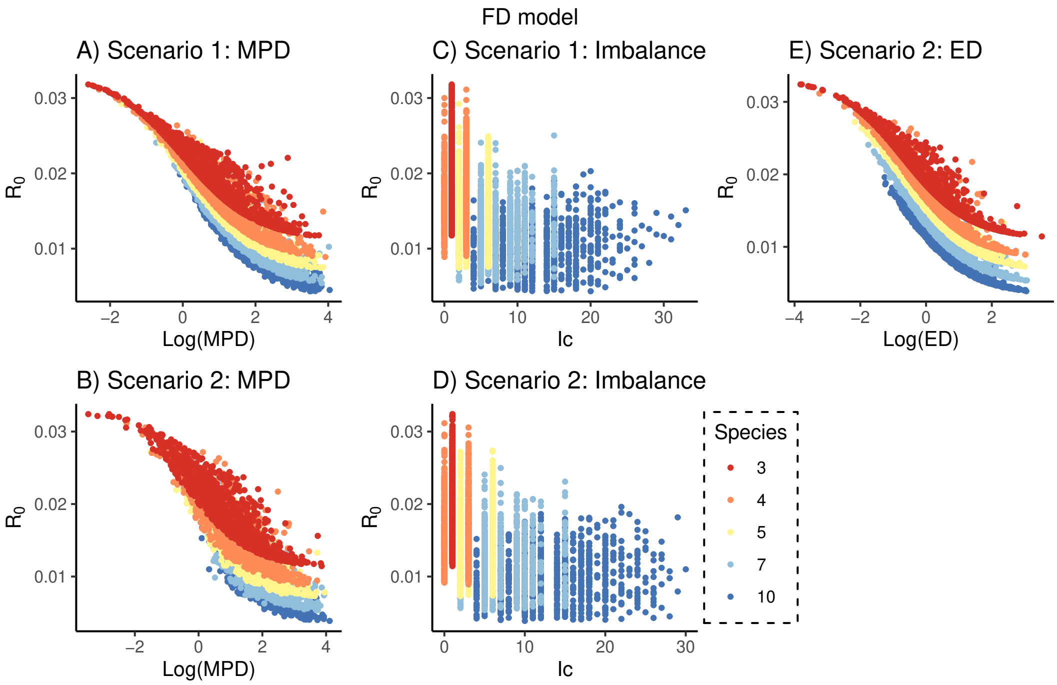


Figure 9: FD model. A,B: Mean Pairwise Distance ($MPD$) of the community is inversely related to $R_0$ in Scenario 1 (A) and Scenario 2 (B). C,D: Phylogenetic Imbalance of the tree connecting host species has variable effects on $R_0$. E: Evolutionary Distinctiveness of the primary host in Scenario 2 decreases $R_0$. Data from simulations with $\psi = 0.01$, $\nu = 0.1$, $\mu = 0.2$. Colours indicate simulated communities with different host richness ($n$=3,4,5,7,10).

### 7 Application to Gilbert *et al.* (2012) functions

Here, we illustrate alternative model parameterized from Gilbert et al. (2012) for different pathogen groups. These functions were derived from plant-pathogen associations as we describe in the main text. Using the general function $logit(c_{ij}) = b_0 - b_1 * log_{10}(PPd_{ij} + 1)$, we used:

For fungi: $b_0$ = 4.40, $b_1$ = -3.32 ($n$ = 95);

For oomycetes: $b_0$ = 2.08, $b_1$ = -2.67 ($n$ = 32);

For insects: $b_0$ = 3.24, $b_1$ = -2.70 ($n$ = 637);

For mites: $b_0$ = 1.96, $b_1$ = -2.17 ($n$ = 87);

For mollusks: $b_0$ = -0.47, $b_1$ = -1.14 ($n$ = 37);

For nematodes: $b_0$ = 2.72, $b_1$ = -2.62 ($n$ = 70);

For plants: $b_0$ = 2.00, $b_1$ = -2.28 ($n$ = 53).

The results for viruses and bacteria are shown in the main text.

Applying the empirical data from the pathogens described above to our model yielded the same results for MPD and Ic as in main text Figure 4. However, there were differences in regressions $R^2$. All pathogen groups show a relatively weak effect of $MPD$ on $R_0$ when the host community is composed of closely related species, allowing frequent disease-sharing (and spillover) between host species. This is most notably the case for fungi, with ectoparasites such as insects, mites, mollusks, plants and nematodes, tending to show slightly greater sensitivity to host phylogenetic structure.

Table 2 shows the results for the multivariate regressions for each of the pathogen groups. Overall, the model for viruses had the lowest explanatory power, which we discuss further in the main text.

| **Pathogen group** | **Model $R^2$** |
|---|---|
| Bacteria | 0.87 |
| Fungi | 0.78 |
| Oomycetes | 0.94 |
| Insects | 0.86 |
| Mites | 0.93 |
| Mollusks | 0.9621 |
| Nematodes | 0.90 |
| Viruses | 0.49 |
| Plants | 0.93 |

Table 2: Multivariate regressions (Scenario 1: $R_0 = MPD + I_c + n$) for the different pathogen groups from the empirical data collected by Gilbert et al. (2012) using model Option 1. All parameter estimates were significant.
